# High-throughput 16S rRNA gene sequencing reveals gut microbial changes in 6-hydroxydopamine-induced Parkinson’s disease mice

**DOI:** 10.1101/633230

**Authors:** Jin Gyu Choi, Eugene Huh, Namkwon Kim, Dong-Hyun Kim, Myung Sook Oh

**Affiliations:** Department of Pharmacy, College of Pharmacy, Kyung Hee University, 26, Kyungheedae-ro, Dongdaemun-gu, Seoul, 02447, Republic of Korea; Department of Medical Science of Meridian, Graduate School, Kyung Hee University, 26 Kyungheedae-ro, Dongdaemun-gu, Seoul, 02447, Republic of Korea; Department of Life and Nanopharmaceutical Sciences, Graduate School, Kyung Hee University, 26, Kyungheedae-ro, Dongdaemun-gu, Seoul, 02447, Republic of Korea; Department of Oriental Pharmaceutical Science, College of Pharmacy and Kyung Hee East-West Pharmaceutical Research Institute, Kyung Hee University, 26, Kyungheedae-ro, Dongdaemun-gu, Seoul, 02447, Republic of Korea

**Keywords:** Parkinson’s disease, gut microbiota, 6-hydroxydopamine, 16S rRNA gene sequencing

## Abstract

Recently, there has been a rapid increase in studies on the relationship between brain diseases and gut microbiota, and clinical evidence on gut microbial changes in Parkinson’s disease (PD) has accumulated. 6-hydroxydopamine (6-OHDA) is a widely used neurotoxin that leads to PD pathogenesis, but whether the alterations of gut microbial community in 6-OHDA-treated mice has not been investigated. Here we performed the 16S rRNA gene sequencing to analyze changes in gut microbial community of mice. We found that there were no significant changes in species richness and its diversity in the 6-OHDA-lesioned mice. The relative abundance of *Lactobacillus gasseri* and *L. reuteri* probiotic species in feces of 6-OHDA-lesioned mice was significantly decreased compared with those of sham-operated mice, while the commensal bacterium *Bacteroides acidifaciens* in 6-OHDA-treated mice was remarkably higher than sham-operated mice. These results provides a baseline for understanding the microbial communities of 6-OHDA-induced PD model to investigate the role of gut microbiota in the pathogenesis of PD.

## Introduction

Parkinson’s disease (PD) is a multicentric neurodegenerative disease clinically defined by motor deficits and progressive degeneration of dopaminergic neurons in brain (1). Non-motor manifestations, which precede the motor disabilities in PD patients, play a key role in the disease progression and evidence for their significance has gradually accumulated (2-4). Among the non-motor symptoms of PD, gastrointestinal (GI) dysfunction, including drooling, impaired gastric emptying, and constipation are frequently reported (5, 6).

Accumulating evidence suggests that the brain is directly involved in gut dysbiosis, an alteration in gut microbiota composition leading to its imbalance, and GI dysfunction following exposure to central stress-like depression (7-9). Gut dysbiosis may cause gut permeability, affecting the GI epithelial barriers and immune system (10, 11). The immune responses triggered by gut microbiota changes could enhance the inflammatory reactions that induce misfolded α-synuclein, which is a pathological hallmark of PD (12, 13).

Scheperjans and his colleagues explored the relationship between gut microbiota changes and clinical phenotypes of PD with fecal microbiome analysis, which showed a reduction in beneficial *Prevotellaceae* and an elevation of pathogenic *Enterobacteriaceae* in PD patients with severe gait disturbance (14). In addition, Keshavarzian et al (15) reported that anti-inflammatory bacterial genera such as *Blautia, Coprococcus*, and *Roseburia* were less abundant in feces of PD patients whereas *Ralstonia* known as a pro-inflammatory bacterial genus was more abundant in the mucosa of PD patients. These studies suggest that changes in gut microbiota composition are closely associated with PD pathogenesis, however, whether gut microbial changes could occur by intracerebral injection of chemical neurotoxins that cause the pathogenesis of PD has not been studied yet.

This study aimed to investigate whether unilateral brain lesions induced by intracerebral injection of 6-hydroxydopamine (6-OHDA) neurotoxin, which not only causes the death of nigrostriatal dopaminergic neurons in brain but also GI dysfunctions such as gastroparesis (16, 17), affects the alteration of gut microbiota composition. To do this, we administrated 6-OHDA directly to the striatum of mouse brain and performed high-throughput sequencing of 16S rRNA genes from fecal samples. Then, we analyzed the alterations of species richness, bacterial diversity, relative abundance at several taxonomic levels, and predicted the functional composition of microbial communities.

## Materials and Methods

### Animals and surgery

Male ICR mice (8 weeks-old) were purchased from Daehan Biolink (Eumseong, Korea). The animals were housed into total 6 cages (3 cages (n=2/cage) and 1 cage (n=3/cage) per sham-operated group; 5 cages (n=2/cage) per 6-OHDA-lesioned group) at an ambient temperature of 23 ± 1°C and relative humidity 60 ± 10% under a 12 h light/dark cycle and were allowed free access to water and food. This study was carried out in accordance with the Principles of Laboratory Animal Care (NIH publication number 80-23, revised 1996). The protocol was approved by the Animal Care and Use Guidelines of Kyung Hee University, Seoul, Korea (Permit number: KHUASP(SE)-16-127). Mice were monitored for total schedule once daily. Mice were euthanized in case of 35% weight loss (humane endpoints) according to the approved protocol, but the euthanized mice were not detected in this study. The unilateral injection of 6-OHDA was performed as modified methods (18). Briefly, mice were anesthetized with tribromoethanol (312.5 mg/kg, *i.p.*) and placed in a stereotaxic apparatus (myNeuroLab, St. Louis, MO, USA). They received a unilateral injection of 2 μl 6-OHDA (16 μg/2 μl; 6-OHDA-lesioned group (n=10)) or equal volume of vehicle (saline with 0.1% ascorbic acid; sham-operated group (n=9)) into the right striatum according to the mouse brain atlas (coordinates with respect to bregma: AP +0.5 mm, ML +2.0 mm, DV: −3.0 mm) (19). After the surgery, mice were maintained body temperature by using heating pads or blankets.

### Fecal sample collection, DNA extraction, and sequencing

Fecal samples were collected at 14 days after the surgery. All samples were placed immediately into sterile plastic tubes and stored at −80°C until analysis. DNA was extracted from the samples using the FastDNA® SPIN Kit for Soil (MP Biomedicals Inc., Solon, USA) according to the manufacturer’s instructions.

PCR amplification was performed using primers targeting from V3 to V4 regions of the 16S rRNA gene with extracted DNA. For bacterial amplification, primers of 341F (5’-TCGTCGGCAGCGTC-AGATGTGTATAAGAGACAG-CCTACGGGNGGCWGCAG-3’; underlining sequence indicates the target region primer) and 805R (5’-GTCTCGTGGGCTCGG-AGATGTGTATAAGAGACAG-GACTACHVGGGTATCTAATCC-3’). The amplifications were carried out under the following conditions: initial denaturation at 95 °C for 3 min, followed by 25 cycles of denaturation at 95 °C for 30 sec, primer annealing at 55 °C for 30 sec, and extension at 72 °C for 30 sec, with a final elongation at 72 °C for 5 min. Then, secondary amplification for attaching the Illumina NexTera barcode was performed with i5 forward primer (5’-AATGATACGGCGACCACCGAGATCTACAC-XXXXXXXX-TCGTCGGCAGCGTC-3’; X indicates the barcode region) and i7 reverse primer (5’-CAAGCAGAAGACGGCATACGAGAT-XXXXXXXX-AGTCTCGTGGGCTCGG-3’).

The condition of secondary amplification is equal to the former one except the amplification cycle set to 8. The PCR product was confirmed by using 2% agarose gel electrophoresis and visualized under a Gel Doc system (BioRad, Hercules, CA, USA). The amplified products were purified with the QIAquick PCR purification kit (Qiagen, Valencia, CA, USA). Equal concentrations of purified products were pooled together and short fragments (non-target products) were removed with an Ampure beads kit (Agencourt Bioscience, MA, USA). The quality and product size were assessed on a Bioanalyzer 2100 (Agilent, Palo Alto, CA, USA) using a DNA 7500 chip. Mixed amplicons were pooled and the sequencing was carried out at ChunLab, Inc. (Seoul, Korea), with an Illumina MiSeq Sequencing system (Illumina, USA) according to the manufacturer’s instructions.

### Taxonomic assignment of sequence reads

Processing raw reads started with quality checking (QC) and filtering of low quality (<Q25) reads by trimmomatic 0.32 (20). After the QC pass, paired-end sequence data were merged together using PandaSeq (21). Primers were then trimmed with ChunLab’s in-house program at a similarity cut off of 0.8. Sequences were denoised using Mothur’s pre-clustering program, which merge sequences and extracts unique sequences allowing up to 2 differences between sequences (22). The EzTaxon database (http://www.eztaxon-e.org/) was used for Taxonomic Assignment using BLAST 2.2.22 and pairwise alignment was used to calculate similarity (23, 24). Microbiome taxonomic profiling was analyzed by BIOiPLUG program (ChunLab Inc., Seoul, Korea). The uchime and non-chimeric 16S rRNA database from EzTaxon were used to detect chimera on reads that contained less than 97% best hit similarity rate (25). Sequence data is then clustered using -Hit and UCLUST, and alpha diversity analysis was carried out (26, 27).

## Statistical analysis

All statistical parameters were calculated using GraphPad Prism 5.0 software (GraphPad Software Inc., San Diego, USA). Values were expressed as the median and quartiles of the data. The results were analyzed with Student’s t-test between two groups. Differences with a *p* value less than 0.05 were deemed to be statistically significant.

Linear discriminant analysis (LDA) effect size (LEfSe) based on Phylogenetic Investigation of Communities by Reconstruction of Unobserved States (PICRUSt) was used to predict how taxonomic differences between fecal microbiota of two groups impact their microbial metabolic potential in the main functional classes (KEGG categories) (28). LEfSe uses the Kruskal–Wallis rank-sum test to identify features with significantly different abundances between assigned taxa compared to the groups, and LDA to estimate the size effect of each feature, based on a p < 0.05 and LDA score > 2.0. The PICRUSt and LEfSe were analyzed by the EzBioCloud database (ChunLab Inc., Seoul, Korea) (29).

## Results

### The alteration of bacterial species richness in 6-OHDA-lesioned mice

All analyzed sequences contained at least 2 of the V3 and V4 16S rRNA gene regions (30). Each read was taxonomically assigned according to the EzTaxon database. When each phylotype at species level was defined using a baseline of 97% nucleotide sequence similarity from 883 to 1,510 phylotypes (average 1,097) were found in the analyzed samples. To investigate the alterations of species richness in feces of 6-OHDA-lesioned mice, we estimated the abundance-based coverage estimators (ACE), Chao1, and the number of operational taxonomic unit (OTU) at species level. The values observed in the sham-operated and 6-OHDA-lesioned mice were comparable (Fig. 1).

**Fig. 1.**
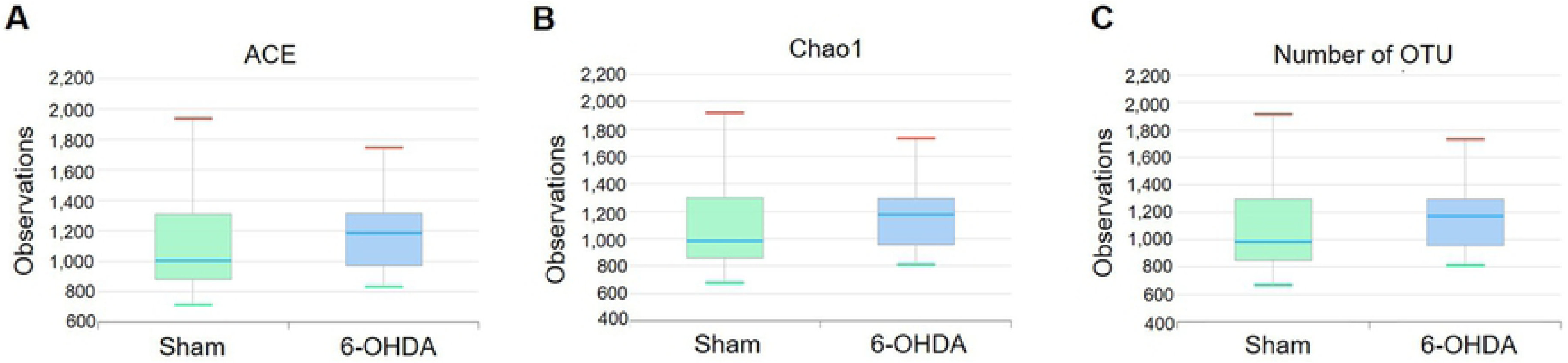
The alteration of bacterial species richness between sham-operated and 6-OHDA-lesioned mice. These data are analyzed by (A) ACE, (B) Chao1, and (C) the number of OTUs. Values are expressed as box and whisker which presents the median and quartiles of the data. n=9/sham-operated group; n=10/6-OHDA-lesioned group.

### The alteration of bacterial diversity in 6-OHDA-lesioned mice

To investigate the alterations of bacterial diversity in mice feces followed by 6-OHDA lesions, we estimated several diversity indexes such as Shannon’s diversity index, Simpson’s diversity index, and phylogenic diversity at species level. We found that there was no difference between the groups (Fig. 2). These data show that bacterial species diversity was unaffected by 6-OHDA lesion.

**Fig. 2.**
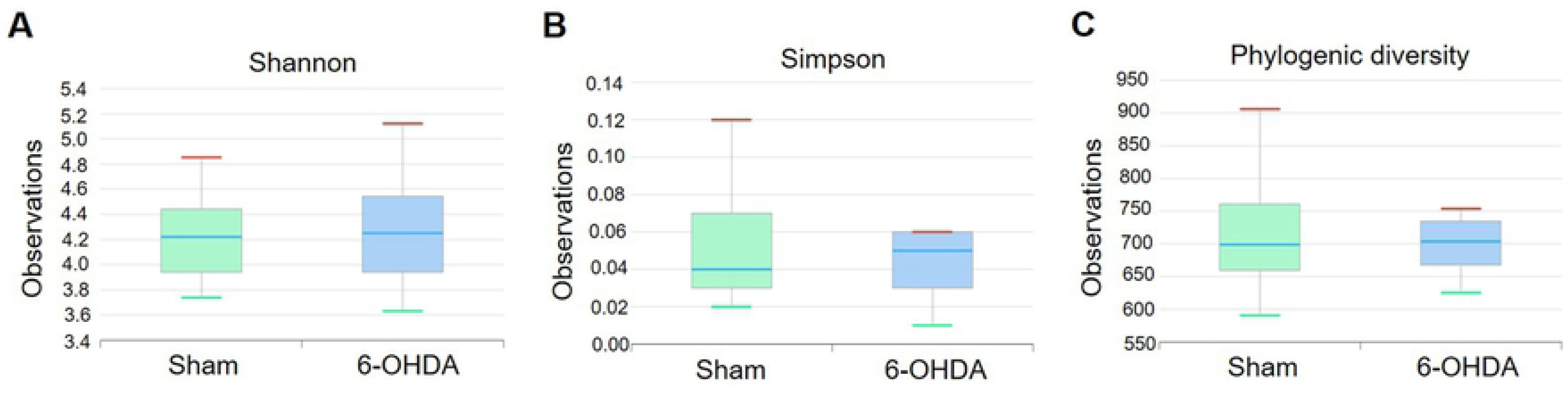
The alteration of bacterial diversity between sham-operated and 6-OHDA-lesioned mice. These data are analyzed by (A) Shannon index, (B) Simpson index, and (C) phylogenic diversity index. Values are expressed as box and whisker which presents the median and quartiles of the data. n=9/sham-operated group; n=10/6-OHDA-lesioned group.

### The alteration of relative abundance in 6-OHDA-lesioned mice

To investigate the alteration of bacterial composition after 6-OHDA administration, the proportion of each taxon at the family, genus, and species levels was compared between sham-operated and 6-OHDA-lesioned groups. The ratio was limited to each taxa containing 1% or more. At the family level, the relative abundance of *Lactobacillaceae* showed a significant decrease in the 6-OHDA-lesioned group (4.07 ± 0.82%) compared with the sham-operated group (13.54 ± 4.65%, p=0.0498), while that of *Bacteroidaceae* was significantly increased in the 6-OHDA-lesioned group (19.32 ± 5.49%) compared with the sham-operated group (6.33 ± 1.94%, p=0.0476). The alteration of other bacterial families, including *S24-7_f, Ruminococcaceae, Prevotellaceae, Lachnospiraceae*, and *Erysipelotrichaceae* was not significant between the groups (Fig 3). The genera *Lactobacillus* was significantly decreased in 6-OHDA-lesioned mice (3.99 ± 0.78%) compared with those of the sham-operated group (13.28 ± 4.39%, p=0.0424), while the *Bacteroides* genus was significantly increased in 6-OHDA-lesioned mice (18.94 ± 4.91%) compared with those of the sham-operated group (6.96 ± 2.18%, p=0.0466). Other genera like *Pseudoflavonifractor, Oscillibacter, Marvinbryantia, Lachnospiraceae LLKB_g, Lachnospiraceae KE159571_g, Lachnospiraceae KE159538_g, Lachnospiraceae JQ084490_g, Muribaculaceae HM124247_g, Muribaculaceae FJ881296_g, Eisenbergiella, Muribaculaceae EF602759_g, Muribaculaceae EF406806_g, Muribaculaceae DQ815871_g, Lachnospiraceae Clostridium_g24, Lachnospiraceae Clostridium_g21, Lachnospiraceae AJ576336_g, Muribaculaceae AB606322_g, S24-7_f_uc, Lachnospiraceae_uc, Prevotella, and Ruminococcaceae JN713389_g* did not show any significant difference between the groups (Fig 4). In Fig. 5, two *Lactobacillus* species like *L. gasseri* and *L. reuteri* were significantly reduced in the 6-OHDA-lesioned group (1.30 ± 0.44% and 0.84 ± 0.18%, respectively) compared to the sham-operated group (6.14 ± 2.22%, p=0.0378 and 3.82 ± 1.37%, p=0.0355, respectively), while the relative abundance of commensal bacteria *B. acidifaciens* was markedly increased in the 6-OHDA-lesioned group (10.85 ± 4.12%) compared with those of the sham-operated group (1.59 ± 0.40%, p=0.0494). There was no significant difference between the groups at the species level of *L. murinus, Clostridium_g24 HM124173_s, Clostridium_g21 HM124119_s, Bacteroides HM124113_s, LLKB_g FJ881271_s, Clostridium_g24 EF604629_s, Bacteroides EF604598_s, DQ815871_g EF603701_s, DQ815871_g EF603109_s, DQ815871_g EF406459_s, Clostridium_g24 EF098800_s, B. vulgatus, AB606322_g AB606322_s, DQ815871_g_uc, KE159538_g KE159628_s, KE159538_g EF605065_s, Prevotella DQ815942_s, KE159538_g DQ815411_s, Eisenbergiella AB626958_s*, and *Lactobacillus_uc*. These results suggest that the remarkable changes of bacterial composition may occur in the mouse intestine after 6-OHDA administration, showing the significant decrease of beneficial bacteria such as *Lactobacillus* probiotic species.

**Fig. 3.**
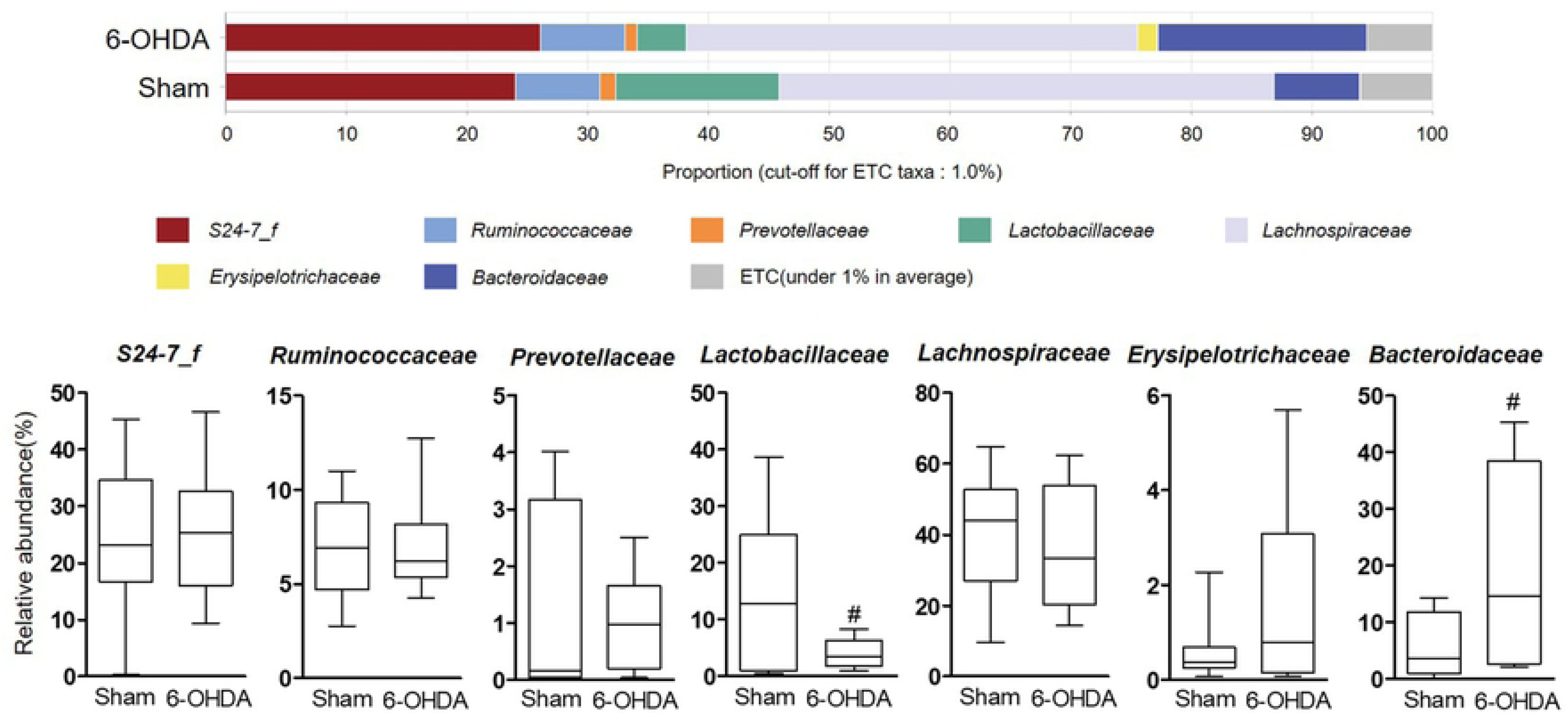
The relative abundance of bacterial family identified in individual samples of sham-operated and 6-OHDA-lesioned mice, respectively. Notable bacterial family over 1% in at least is marked. Values are expressed as box and whisker which presents the median and quartiles of the data. *^#^p*<0.05 compared with the sham-operated group. n=9/sham-operated group; n=10/6-OHDA-lesioned group.

**Fig. 4.**
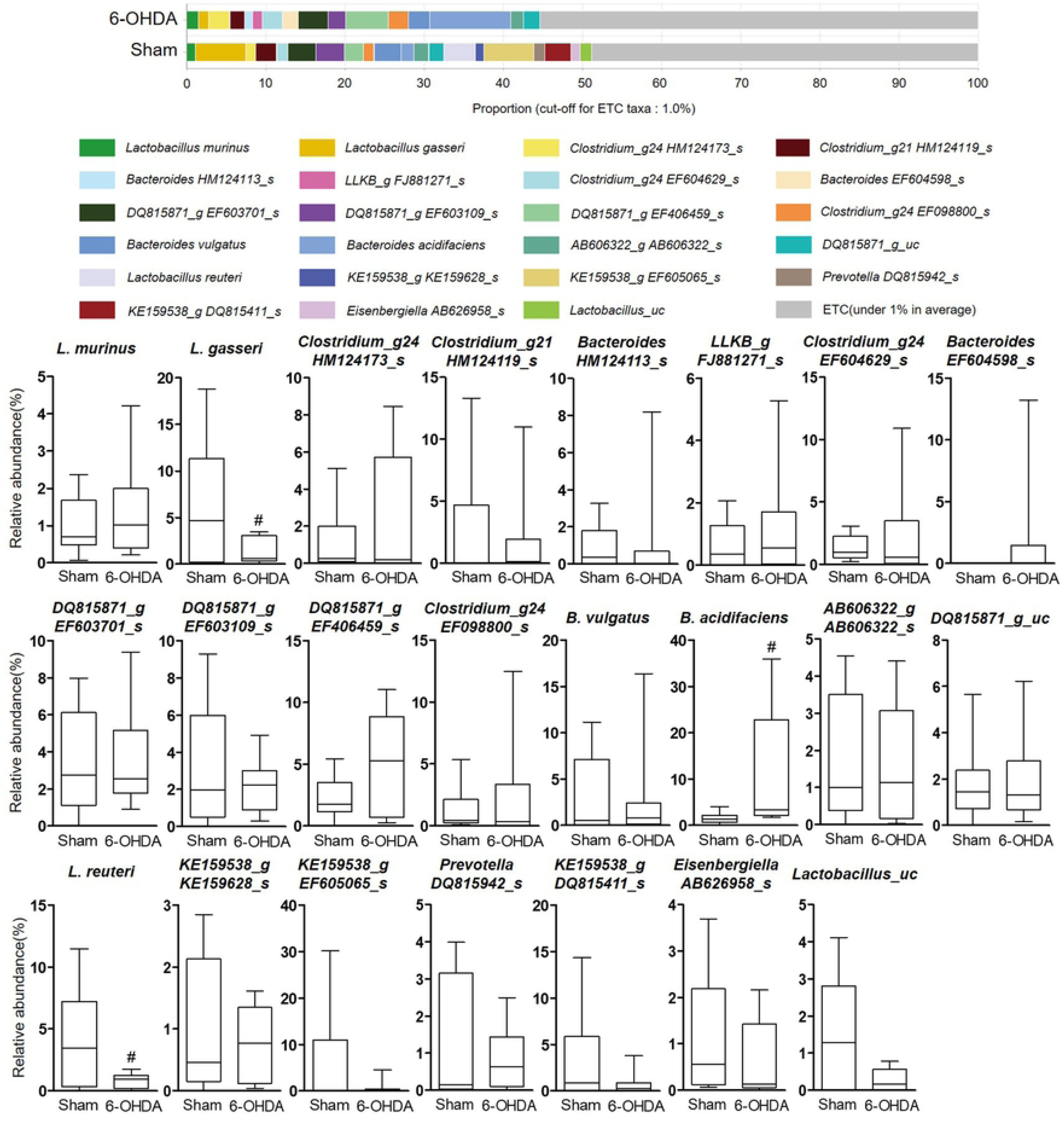
The relative abundance of bacterial genus identified in individual samples of sham-operated and 6-OHDA-lesioned mice, respectively. Notable bacterial genus over 1% in at least is marked. Values are expressed as box and whisker which presents the median and quartiles of the data. *^#^p*<0.05 compared with the sham-operated group. n=9/sham-operated group; n=10/6-OHDA-lesioned group.

**Fig. 5.**
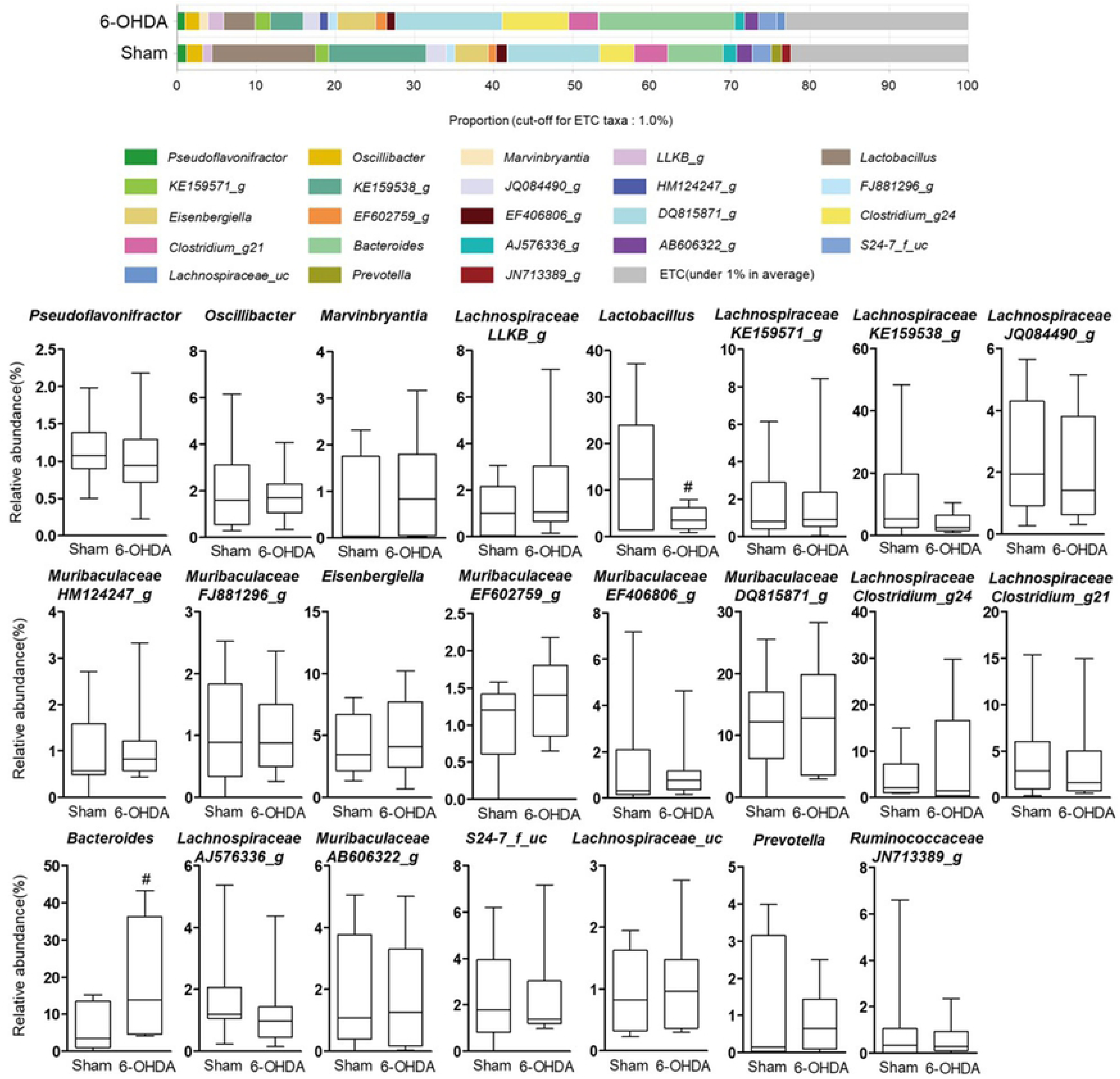
The relative abundance of bacterial species identified in individual samples of sham-operated and 6-OHDA-lesioned mice, respectively. Notable bacterial species over 1% in at least is marked. Values are expressed as box and whisker which presents the median and quartiles of the data. *^#^p*<0.05 compared with the sham-operated group. n=9/sham-operated group; n=10/6-OHDA-lesioned group.

### Predicted functional composition using LEfSe analysis based on the PICRUSt dataset

Within the main KEGG categories, LEfSe analysis indicated that homologous recombination; galactose metabolism; other glycan degradation; starch and sucrose metabolism; amino sugar and nucleotide sugar metabolism; biosynthesis of antibiotics; biosynthesis of amino acids; biosynthesis of secondary metabolites; and metabolic pathways were enriched in 6-OHDA-lesioned group, whereas ribosome; carbon metabolism; quorum sensing; tryptophan metabolism; alanine, aspartate and glutamate metabolism; hepatitis C; ABC transporters metabolism pathway genes were enriched in sham-operated group (Fig 6).

**Fig. 6.**
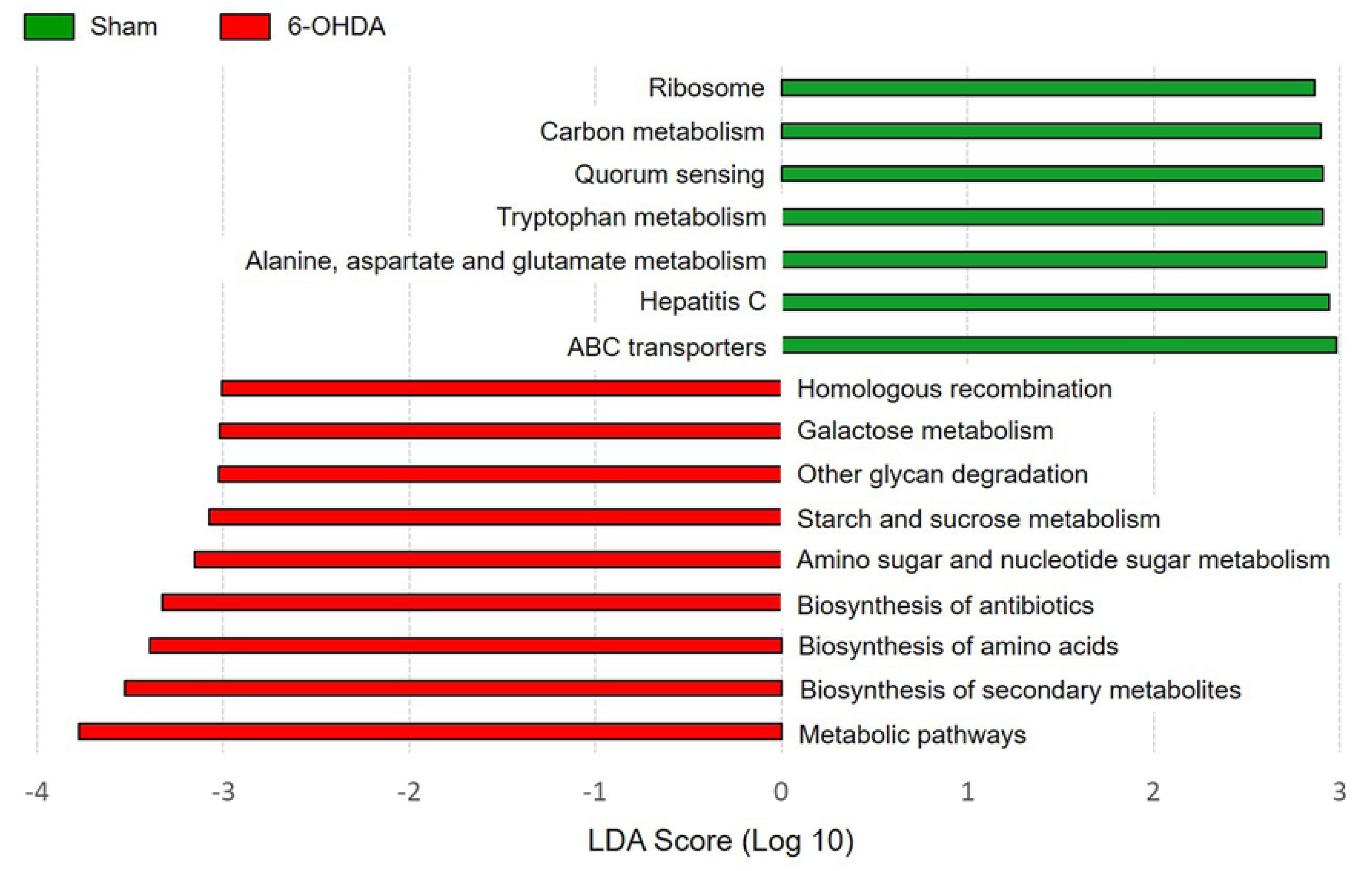
Predicted functional composition of metagenomes based on 16S rRNA gene sequencing data. LEfSE analysis performed on metabolic functions based on PICRUSt results shows several significant differences of KEGG categories between sham and 6-OHDA group. The threshold for the logarithmic LDA score was 2.0 and p < 0.05 for the factorial Kruskal-Wallis test among classes.

## Discussion

The composition of gut microbiota can have a remarkable impact on disease status as well as normal physiology in brain (31, 32). Emerging evidence supports that gut dysbiosis by altered gut microbiota has the potential to be closely linked with neurodegenerative diseases, including PD (33-36). In this study, we found the changes in gut microbiota composition in a PD mouse model by unilateral 6-OHDA-lesion using the high-throughput 16S rRNA gene sequencing method.

First, we confirmed the well-established PD mouse model induced by intracerebral injection of 6-OHDA as shown in severe motor deficits and dopaminergic neuronal damage (Fig S1). In this model, we found little difference between the sham-operated and 6-OHDA-lesioned groups on species richness, bacterial diversity (Fig 1, 2). We also found no significance between two groups on a principal coordinates analysis evaluated by Permutational multivariate analysis of variance (p=0.153, Fig S2). It has been reported that microbial species richness and its diversity indicate whether gut microbiota impacts biological entities, but the ecological meaning of this index simply shows the number of bacterial species, and is not an indicator of gut dysbiosis (37-39).

Next, we observed a significant differentiation in gut microbiota composition at the phylogenic family, genus, and species levels between the sham-operated and 6-OHDA-lesioned groups (Fig. 3-5). We showed a relative lower abundance of *Lactobacillaceae* and *Lactobacillus* at the family and genus level, respectively, thereby the relative abundance of *L. gasseri* and *L. reuteri* was significantly reduced in the 6-OHDA-lesioned group compared to the sham-operated group. Several studies showed that exposure to excessive stress induced a shift in microbial composition that was the reduced proportion of anti-inflammatory bacteria, including *Lactobacillus* (40, 41). These bacteria regulated emotional behavior by inducing transcription of γ-aminobutyric acid receptors and suppressed the disease progression and excessive T cells-mediated immune responses in experimental autoimmune encephalomyelitis-induced mouse model of multiple sclerosis (42-44); especially *L. reuteri* supplementation restored gut motility in patients with chronic constipation, which is a major non-motor symptoms in PD (45). We found a clear difference in microbial community patterns between the sham-operated and 6-OHDA-lesioned groups. Several reports showed that *Lactobacillus* was more abundant in advanced PD patients than controls on the contrary to our results (14, 46-48). This implies that *Lactobacillus* abundance may be different by various factors. We also found the remarkable increase of *Bacteroides* abundance after the 6-OHDA lesion. Keshavarzian and his colleagues reported that the counts of *Bacteroides* were significantly increased in fecal samples of PD patients (15). This remarkable increase is closely related to the immunoglobulin A (IgA), which has been increased in mice exposure to repeated 6-OHDA or inoculation of germ-free mice with *B. acidifaciens* (49, 50). It is reported that IgA plays a critical role in maintenance of gut microbiota composition and is presumed to prevent intestinal damages from external pathogen (51, 52). Thus, both the relative abundance of *Lactobacillus* and *Bacteroides* in 6-OHDA-lesioned mice suggest the imbalanced gut microbiota composition status seen in intestinal pathological condition by external toxin, including 6-OHDA.

In a recent microbiome analysis of the cecal sample in mice induced by intraperitoneal injection of 6-OHDA, peripheral 6-OHDA induced an approximately 20-fold decrease in the *Prevotellaceae* compared with control group (53). This implies that peripheral 6-OHDA, which does not cross the blood-brain barrier (54), can directly affect enteric microbiota. Our results showed the different patterns of gut microbial changes unlike those of peripheral 6-OHDA injection. It is expected that intracerebral injection of 6-OHDA induced intestinal microbial changes through indirect regulation which can affect these changes. Recent studies indicate that activation of the hypothalamic-pituitary-adrenal (HPA) axis following exposure to chronic or acute stress affects gut microbiota composition directly via stimulation of the vagus nerve (40, 55, 56). Moreover, the release of neurotransmitters in the brain through cholinergic sympathetic activation influences changes in gut microbiota composition and GI functions (57). It is possible that brain damage induced by 6-OHDA, which can regulate the HPA axis and eliminate cholinergic sympathetic innervation (58, 59), may interfere with the signaling from brain to gut along the HPA axis, thereby alters gut microbiota composition.

LEfSe analysis identified several categories of functional metabolism that were enriched in sham and 6-OHDA mice. Mice with 6-OHDA lesion had increased abundances of genes responsible for carbohydrate metabolism (eg. galactose, starch, sucrose and glycan degradation), homologous recombination, biosynthesis of antibiotics, and biosynthesis of secondary metabolites. Especially, alterations in secondary metabolites have been observed in the gut microbiota metagenomics data of PD patients, suggesting that metabolic dysfunction of secondary metabolites may be an important feature of PD (60). However, this result is only originated from metagenomic functions and metabolomic approaches are preferred to identify factual changes in metabolic function of microbiota of 6-OHDA-lesioned mice.

There are several limitations to this study. First, analysis of the mouse gut microbiota was performed fourteen days after 6-OHDA injection. The time-course study after intracerebral injection of 6-OHDA on gut microbial changes would provide more detailed information. Second, how 6-OHDA could induce changes of gut microbiota composition in mice was unexplored, then further study should focus on the mechanism of 6-OHDA-induced gut microbial changes. We first identified the relative abundances at taxonomic levels and predicted functional metabolic pathways between sham-operated and 6-OHDA-lesioned groups by the analysis of bacterial 16S rRNA gene sequencing. This study demonstrated that the levels of some microorganisms in the feces, such as *Lactobacillus* and *Bacteroides*, are associated with the exposure to 6-OHDA. These results provides a baseline for understanding the microbial communities of 6-OHDA-induced PD model to investigate the role of gut microbiota in the pathogenesis of PD.

## Authors contributions

**Conceptualization:** JGC, MSO.

**Data Curation:** JGC, NK.

**Formal Analysis:** NK, EH.

**Funding Acquisition:** MSO.

**Investigation:** JGC, EH, NK.

**Methodology:** EH.

**Project Administration:** JGC, MSO.

**Resources:** NK, EH.

**Supervision:** DHK, MSO.

**Validation:** EH, NK.

**Visualization:** JGC.

**Writing – Original Draft Preparation:** JGC.

**Writing – Review & Editing:** DHK, MSO.

